# Assumptions about frequency-dependent architectures of complex traits bias measures of functional enrichment

**DOI:** 10.1101/2020.10.23.352427

**Authors:** Shadi Zabad, Aaron P. Ragsdale, Rosie Sun, Yue Li, Simon Gravel

## Abstract

Linkage-Disequilibrium Score Regression (LDSC) is a popular framework for analyzing GWAS summary statistics that allows for estimating SNP heritability, confounding, and functional enrichment of genetic variants with different annotations. Recent work has highlighted the influence of implicit and explicit assumptions of the model on the biological interpretation of the results. In this work, we explored a formulation of LDSC that replaces the *r*^2^ measure of LD with a recently-proposed unbiased estimator of the *D*^2^ statistic. In addition to modest statistical difference across estimators, this derivation highlighted implicit and unrealistic assumptions about the relationship between allele frequency, effect size, and annotation status. We carry out a systematic comparison of alternative LDSC formulations by applying them to summary statistics from 47 GWAS traits. Our results show that commonly used models likely underestimate functional enrichment. These results highlight the importance of calibrating the LDSC model to achieve a more robust understanding of polygenic traits.

## 1 Introduction

Linkage-Disequilibrium Score Regression (LDSC) provides a general framework for understanding the architecture of polygenic traits from GWAS summary statistics (Bulik-Sullivan, Loh, et al., 2015; Pasaniuc and Price, 2017). Because of its tractability and computational efficiency, it has become a central tool to estimate statistical and population genetic quantities, such as SNP heritability 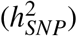 and confounding in GWAS results (Bulik-Sullivan, Loh, et al., 2015), functional enrichment (Finucane et al., 2015), cross-trait genetic correlations (Bulik-Sullivan, Finucane, et al., 2015) as well as genetic correlations across ethnic groups for the same trait (Shi et al., 2019). This diversity of applications stems from the ease of incorporating various assumptions about the architecture of a trait in the LDSC framework. In its original formulation, the LDSC model made the implicit assumption that the expected phenotypic variance explained by each genetic variant was independent of allele frequency. However, subsequent work has shown that this assumption is restrictive and results in downwardly biased estimates of heritability (Speed, Cai, Johnson, Nejentsev, and Balding, 2017).

To guard against these potential sources of bias, the trend in the field has been towards incorporating more expressive models that describe the expected squared effect of a variant as a function of different genetic annotations (Finucane et al., 2015; Gazal et al., 2017; Gazal et al., 2018; Hormozdiari et al., 2018; Hujoel, Gazal, Hormozdiari, van de Geijn, and Price, 2019), such as MAF decile bins and whether a SNP occurs in an annotated enhancer region. An advantage of this approach, commonly referred to as Stratified LDSC (S-LDSC), is that it allows for arbitrary relationships between allele frequency and mean squared effect sizes, thus reducing the need for a priori assumptions that can bias results. It also provides measures of functional enrichment for the categories incorporated into the model. Despite these many advantages, previous work has shown that the way we model the dependence of effect size on functional annotations can result in substantially different estimates of partitioned heritability and functional enrichment (Gazal, Marquez-Luna, Finucane, and Price, 2019; Speed et al., 2017; Speed, Hemani, Johnson, and Balding, 2012).

In this work, we reformulate the LD Score regression framework by replacing the *r*^2^ measure of LD with an estimate based on the *D*^2^ statistic, which has desirable statistical properties (Ragsdale and Gravel, 2020). Even though the choice of the LD statistic itself has little effect on heritability estimates, this formulation highlights implicit assumptions of commonly-used stratified models. Specifically, these models imply relationships between allele frequency, effect size, and annotation status that do not agree with empirical observations. While these assumptions have a small impact on global estimates of SNP heritability, we show that they can result in systematic biases in estimates of functional enrichment. To correct for these biases, we propose using modified S-LDSC models that better capture the empirical relationship between allele frequency and effect size for annotated variants.

## 2 Materials and Methods

### 2.1 LD Score Regression as a function of *D*^2^

In our formulation of LD Score Regression, we assume a standard linear polygenic model *Y* = *Xβ* + *ε*, where *Y* is a vector of phenotypes for *N* individuals, *X* is a *N* × *M* mean-centered (but not variance-normalized) matrix that encodes the genotype at *M* SNPs, *β* is a vector of random and independent effect sizes, and *ε* is a vector of random and independent environmental and other effects. Under this model, assuming no confounding, the expectation of the *χ*^2^ association statistic at locus *j* can be expressed as a function of the squared covariance in allele frequency 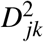 at variants *j* and *k* as:

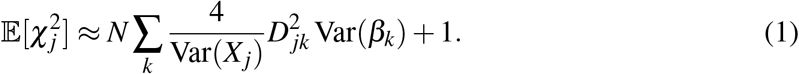

Here, Var(*β*_*k*_) is the variance of causal effect size at SNP *k* and *Var*(*X*_*j*_) = 2*p*_*j*_(1 − *p* _*j*_) is the variance in genotype across individuals at the focal SNP *j*, where *p* _*j*_ is its minor allele frequency in the population. In LDSC, we seek to build a parameterized model for Var(*β*_*k*_) that can be fitted to the observed *χ*^2^. Since Var(*β*_*k*_) is the mean squared effect size for variant *k*, we can then sum over all SNPs to obtain an estimate of SNP heritability. While the simplest model takes Var(*β*_*k*_) to be a constant across SNPs, various models have been proposed that take into account allele frequencies as well as functional annotations. Here we consider models of the form

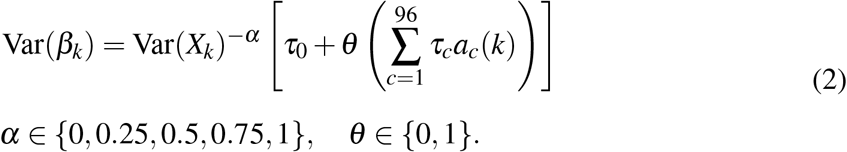

The mean squared effect size Var(*β*_*k*_) of SNP *k* is defined in terms of two components: a dependence on minor allele frequency (MAF) through the parameter *α*, and a dependence on a set of 96 functional annotations *a*_*c*_(*k*) that were incorporated into the baseline-LD model (v2.2) (Gazal et al., 2017; Gazal et al., 2019, see Web Resources). This dependence is controlled by the parameter *θ*. If *θ* = 0, we recover the univariate models introduced in Bulik-Sullivan, Loh, et al., 2015. If *θ* = 1 we obtain the stratified models that are most commonly used in more recent work (Finucane et al., 2015; Gazal et al., 2017; Gazal et al., 2018; Hormozdiari et al., 2018; Hujoel et al., 2019).

The contribution of each annotation *a*_*c*_ to SNP heritability is modulated by the corresponding parameter *τ*_*c*_, which is estimated in the LDSC regression procedure. In this formulation, *τ*_0_ corresponds to the “base” annotation that includes all SNPs (e.g. *a*_0_(*k*) = 1 for all *k*). Overall, in this framework, the expected heritability contributed by a SNP is defined as 𝔼 [*h*_*j*_] = *Var*(*X*_*j*_)*Var*(*β*_*j*_) (Speed and Balding, 2019; Speed, Holmes, and Balding, 2020). Table 1 summarizes some of the relevant models that will be analyzed in this paper.

**Table 1:** The SNP-heritability models analyzed in this paper.

| Heritability model | Parameters | Comments |
| --- | --- | --- |
| Constant effect size model | $\alpha = 0; \theta = 0$ | The variance in effect size $\text{Var}(\beta_j)$ is independent of allele frequency. |
| GCTA model (Yang et al., 2010, Bulik-Sullivan, Loh, et al., 2015) | $\alpha = 1; \theta = 0$ | The expected heritability $\mathbb{E}[h_j]$ contributed by a variant is constant. |
| Baseline-LD model (v2.2) (Gazal, Marquez-Luna, Finucane, and Price, 2019) | $\alpha = 1; \theta = 1$ | A stratified LDSC model that includes a total of 96 functional annotations, including continuous LD-related annotations and MAF bins. This model gives more weight to annotations found in rare variants. |
| MAF-independent baseline-LD model | $\alpha = 0; \theta = 1$ | Similar to the baseline-LD model, though the effect of each annotation is independent of allele frequency. |

In previous work, *α* has often been treated as a fixed parameter, usually set to *α* = 1. In the univariate case (*θ* = 0), this is a mathematically convenient choice that implies constant variance explained by all SNPs and leads to simple expressions of LD scores in terms of the LD statistic *r*^2^ (as detailed below). In turn, this implies that rare variants have much larger per-allele biological effect than common variants. In other research settings, *α* is treated as a continuous parameter that can be fit to data to measure the strength of negative selection for a given trait. Average inferred values of *α* for a number of UK Biobank traits ranged from 0.2 to 0.5, with a mean value of roughly 0.38 (Schoech et al., 2019; Speed et al., 2017; Zeng et al., 2018). However, these values of *α* were inferred for the univariate setting and it is not clear if the interaction between MAF and the various functional annotations can still be characterized by these global estimates. In fact, a recent report using an updated version of the LDAK and SumHer models (Speed and Balding, 2019; Speed et al., 2012; Speed et al., 2020) found variations in the *α* values ranging from 0.23 to 0.67 for SNPs in all but one functional category, and 0.25 for all variants.

Despite these observations, in standard applications of the stratified LDSC models (*θ* = 1), it is still commonly assumed that *α* = 1 (Gazal et al., 2017; Gazal et al., 2018; Hormozdiari et al., 2018; Hujoel et al., 2019). The choice *α* = 1 is not particularly biologically plausible — it is a leftover from the mathematically convenient formulation of LDSC in terms of the *r*^2^ statistic. Equation (2) shows that the assumption *α* = 1 in the stratified setting implies that the contribution of each annotation to predicted effect size is much larger for rare variants. Put another way, under the *α* = 1 model, rare variants in a given functionally enriched category *c* are predicted to have much larger per-allele biological effect than common variants in the same category. Since this assumption has been found to not be the valid in genome-wide analyses (Schoech et al., 2019; Zeng et al., 2018), there is no compelling reason to think that it holds within specific categories of variants.

## 3 Results

### 3.1 Comparing LD scores across models and estimators

The family of models outlined in Equations (1) and (2) may be equivalently expressed in terms of LD scores:

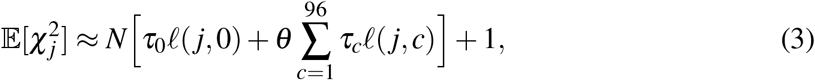

where the LD score *𝓁* (*j, c*) of variant *j* and category *c* is defined as the sum of the annotation value *a*_*c*_(*k*) multiplied by a MAF-weighted linkage disequilibrium term *L*_*jk*_(*α*) over all neighbouring SNPs *k*:

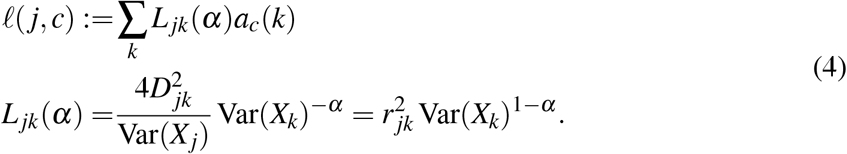

In standard applications of LDSC, the MAF-weighted LD measure *L*_*jk*_(*α*) is typically taken to be 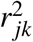, which implies *α* = 1. However, for arbitrary *α*, it can be expressed more generally in terms of a product of the variances at SNPs *j* and *k* with the squared covariance 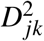 or the squared correlation 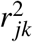. This generalized measure can be estimated in-sample or from a reference panel using, e.g, the following two estimators:

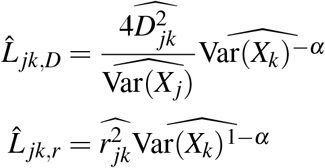

where 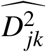 is the unbiased estimator of Ragsdale and Gravel, 2020 and 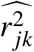 is the bias-corrected estimator used in the original LDSC model (Bulik-Sullivan, Loh, et al., 2015; Yin and Fan, 2001). The estimator for 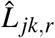 has a particularly simple form for *α* = 1, which is probably a reason for the popularity of this parameter choice. To evaluate whether choices of estimators affect LDSC regression results, we analyzed the differences between the two statistical estimators across different values of *α*. To perform this analysis, we computed LD scores according to Equation (4) using both estimators and *α* values of {0, 0.25, 0.5, 0.75, 1} for three of the super populations in the 1000 Genomes Project (Africans, Asians and Europeans) (Altshuler et al., 2010; The 1000 Genomes Project Consortium, 2012).

Both estimators produce highly correlated estimates (*R*^2^ = 0.99) (Figure 1 (a)), and their bias is comparable even for small panel sample size (Supplementary Material, Supplementary Figure 1). The consistency of these two estimators suggests that they perform equally well in estimating LD scores irrespective of the value of *α*. Thus, biological realism might be a more relevant criterion when choosing *α* than statistical convenience. For the remainder of the discussion in the main text, we will use the 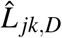 estimator.

**Figure 1.**
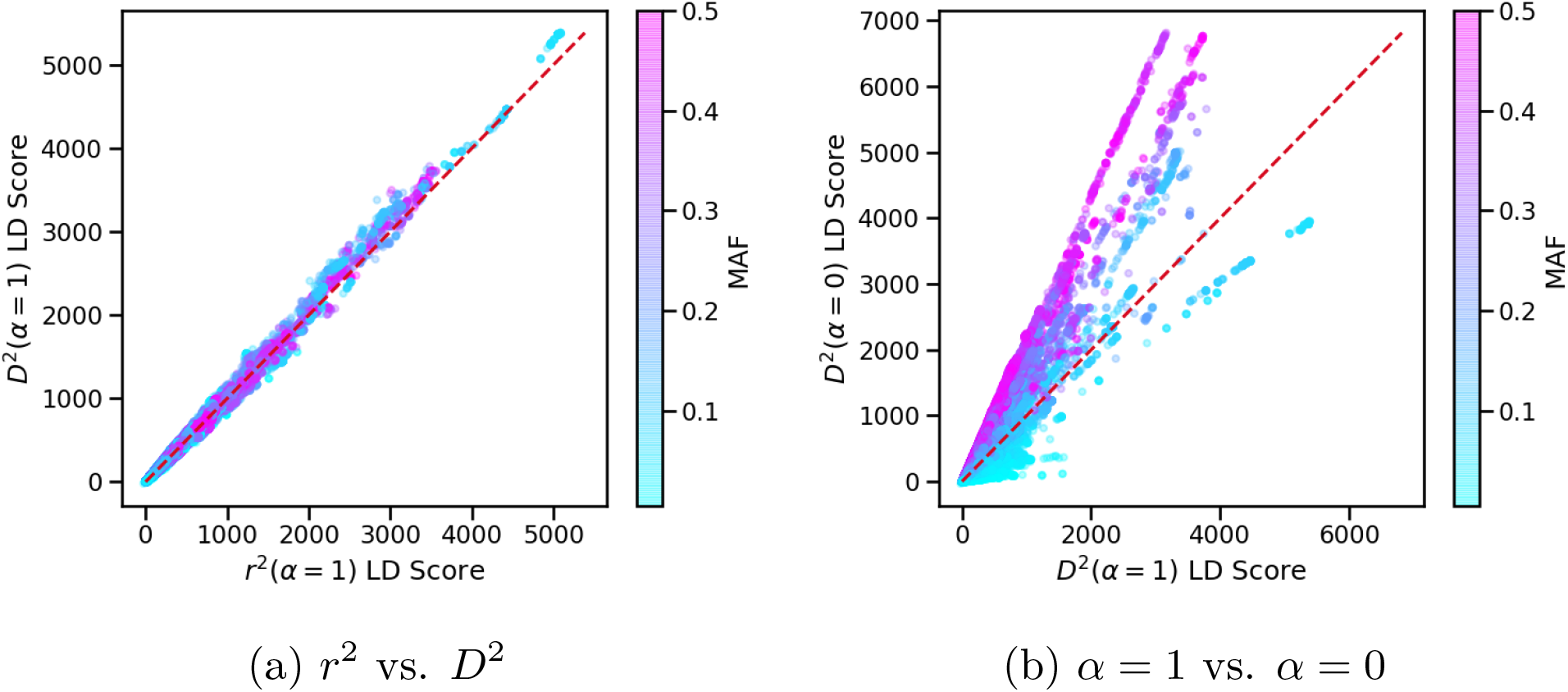
Genome-wide comparison of LD Scores in the European samples (*N* = 489) in the 1000 Genomes Project. **(a)** shows the distribution of LD Scores obtained from the *r*^2^ (x-axis) and *D*^2^ (y-axis) estimators. In **(b)** we show the normalized LD Scores from the *D*^2^ estimator but computed according to different models of SNP heritability (*α* = 1 on the x-axis and the *α* = 0 on the y-axis). Points are colored by the focal SNP’s minor allele frequency (MAF).

Next, we sought to highlight the influence of model assumptions on the LD scores computed from the 1000 Genomes data. For clarity of exposition, we focus on the univariate case where the LDSC model can be written as: 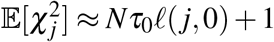 and the LD score is simply the sum of the generalized LD measures *L*_*jk*_(*α*). To qualitatively compare the LD scores with different values of *α* in a consistent manner, we normalize them such that the slope of the univariate regression becomes 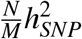, where 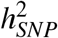 is the total heritability (see Appendix C.1).

As expected, Figure 1(b) shows that common variants have higher LD scores under the *α* = 0 model compared to the *α* = 1 model, and the opposite trend is observed for rare variants. This effect is less stark for intermediate values of *α* (Supplementary Figure 2). In practical terms, the choice of *α* determines which categories of SNPs will have higher weight in the LDSC regression and, as has been documented before (Speed et al., 2017), this will in turn influence the global estimates of SNP heritability.

**Figure 2.**
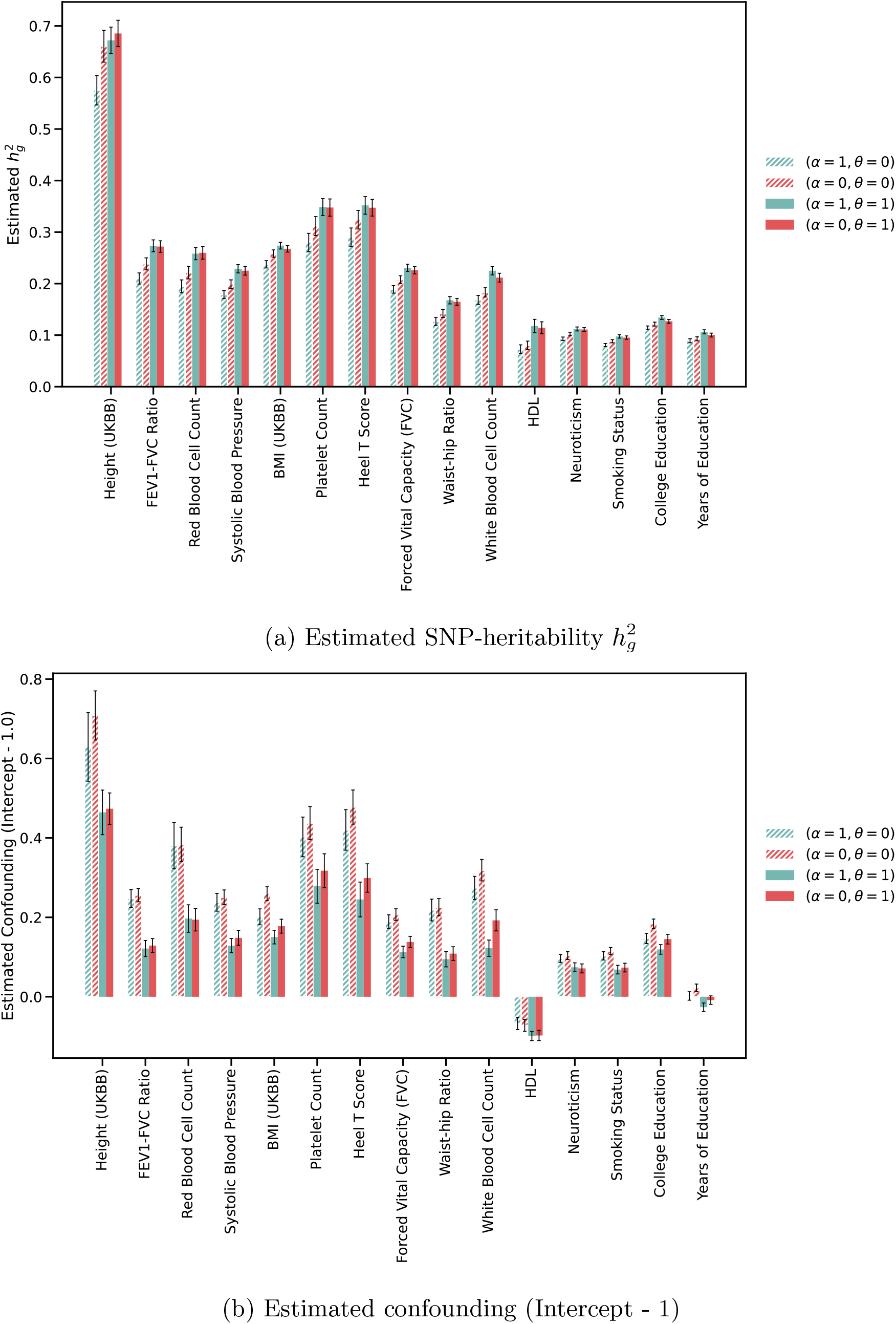
Estimates of SNP heritability and confounding for 15 GWAS traits using 4 different models of heritability. Dashed bars show estimates for univariate models. Solid bars show estimates for stratified models. Error bars correspond to jackknife standard errors. Color code: turquoise (*α* = 1), red (*α* = 0).

### 3.2 Estimates of SNP-heritability and Confounding for 47 GWAS Traits

To understand the implications of choosing different values of *α* on global estimates of SNP heritability and confounding, we applied the LD Score regression framework to a total of 47 GWAS traits (41 independent traits), for which summary statistics have been previously analyzed (Gazal et al., 2019; Hujoel et al., 2019) (Supplementary Table S1, see Web Resources). We confirmed that the choice of statistical estimator has a modest impact on estimates of global heritability and confounding (Supplementary Table S7): for example, the 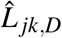 estimator in the *α* = 1 stratified model gave lower estimates of heritability than the 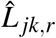 estimator for all 47 traits, but the relative differences were always smaller than 6% (Supplementary Table S3), and always smaller than twice the block-jackknife standard error of the estimator. By contrast, the choice of *α* can produce large differences.

For the univariate models (*θ* = 0), we find that estimates of heritability with *α* = 0 are different by more than two standard errors to the *α* = 1 estimate for 29 out of the 47 traits considered (Figure 2, Supplementary Figure 6, Supplementary Tables S5-6). In general, we observe that the univariate models with *α* = 0 produce heritability estimates that are closer to the estimates from the stratified models for 44 out of 47 traits (Figure 2(a), Supplementary Tables S2-4), which is consistent with previous estimates of *α* values that are closer to zero (*α* ≈ 0.38, Schoech et al., 2019; Zeng et al., 2018). This trend is reversed for the intercept, with the *α* = 1 model producing estimates that are closer to the stratified models for most traits. Supplementary Figure 6(b) shows that intermediate values of *α*, such as *α* = 0.25, produce estimates of confounding and global heritability that are closer to the stratified models than both *α* = 0 and *α* = 1.

In the case of the stratified models (*θ* = 1), the differences are more subtle. Out of the 47 traits analyzed, none showed a significant difference in the global estimates of SNP heritability across the models with *α* = 0 and *α* = 1 (Supplementary Tables S2-4). At the same time, small but significant differences are observed in the estimates of the intercept for 3 of the traits analyzed (Eosinophil Count, Tanning, and White Blood Cell Count) (Supplementary Tables S2-4). Overall, our analysis confirms that the stratified models (with the MAF decile bins included) successfully counteract the bias induced by an arbitrary choice of *α*, producing global estimates of SNP heritability and confounding that are largely concordant across different choices of *α* (Figure 2, Supplementary Figure 7).

### 3.3 Examining Estimates of Coefficients and Functional Enrichment

The preceding analyses showed that the stratified models generally produce concordant estimates of global parameters such as SNP heritability and confounding, with the choice of *α* having only a minor impact. However, as has been noted above, different choices of *α* can still result in different estimates for quantities associated with partitioned heritability, such as standardized heritability coefficients and functional enrichment. To examine the influence of *α* on these partitioned heritability metrics, we first focus on the per-standardized annotation coefficients, as defined by Gazal et al., 2017:

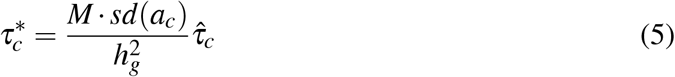

The quantity 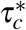 is defined as additive change in per-SNP heritability associated to a 1 standard deviation increase in the value of the annotation (*sd*(*a*_*c*_)), normalized by the average per-SNP heritability over all SNPs for the trait 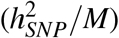. Despite its usefulness for examining contributions to heritability across traits, the *τ*^*^ metric as defined above is not suitable for comparing models with different values of *α*, since the 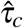 in these models are on different scales (see Appendix B). Here, we propose a modified metric that captures the contributions of the different annotations to heritability as well as the influence of *α*:

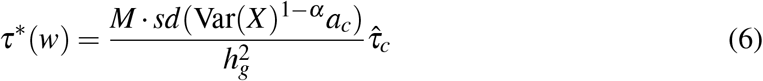

The modified metric *τ*^*^(*w*) has the same overall interpretation as the metric proposed by Gazal et al., 2017, with the main difference being that we multiply the value of the annotation by the variance in allele frequency Var(*X*)^1−*α*^ when computing its standard deviation.

Figure 3(a) shows the estimates for the standardized coefficients for all the functional annotations that we analyzed, meta-analyzed across the 47 GWAS traits. We find that, in general, models with *α* = 0 and *α* = 1 produce comparable estimates of *τ*^*^(*w*), with some notable exceptions (Supplementary Table S8). In particular, the models tend to diverge in the significance they assign to annotations associated with QTLs (e.g. the set of MaxCPP annotations introduced in Hormozdiari et al., 2018, with the *α* = 0 models often failing to reach the Bonferroni significance threshold for those annotations (3(a), Supplementary Table S8). Comparable differences between the models are also observed for some of the LD-related annotations (e.g. CpG Content and Recombination Rate) that were introduced by Gazal et al., 2017 (Figure 3(c), Supplementary Table S8). Note that some of these differences persist for intermediate values of *α*, though at a much smaller scale (Supplementary Figures 8-9). Thus the choice of *α* can influence the predicted biological relevance of different annotations even when frequency bins are used to model frequency-dependent effects.

**Figure 3.**
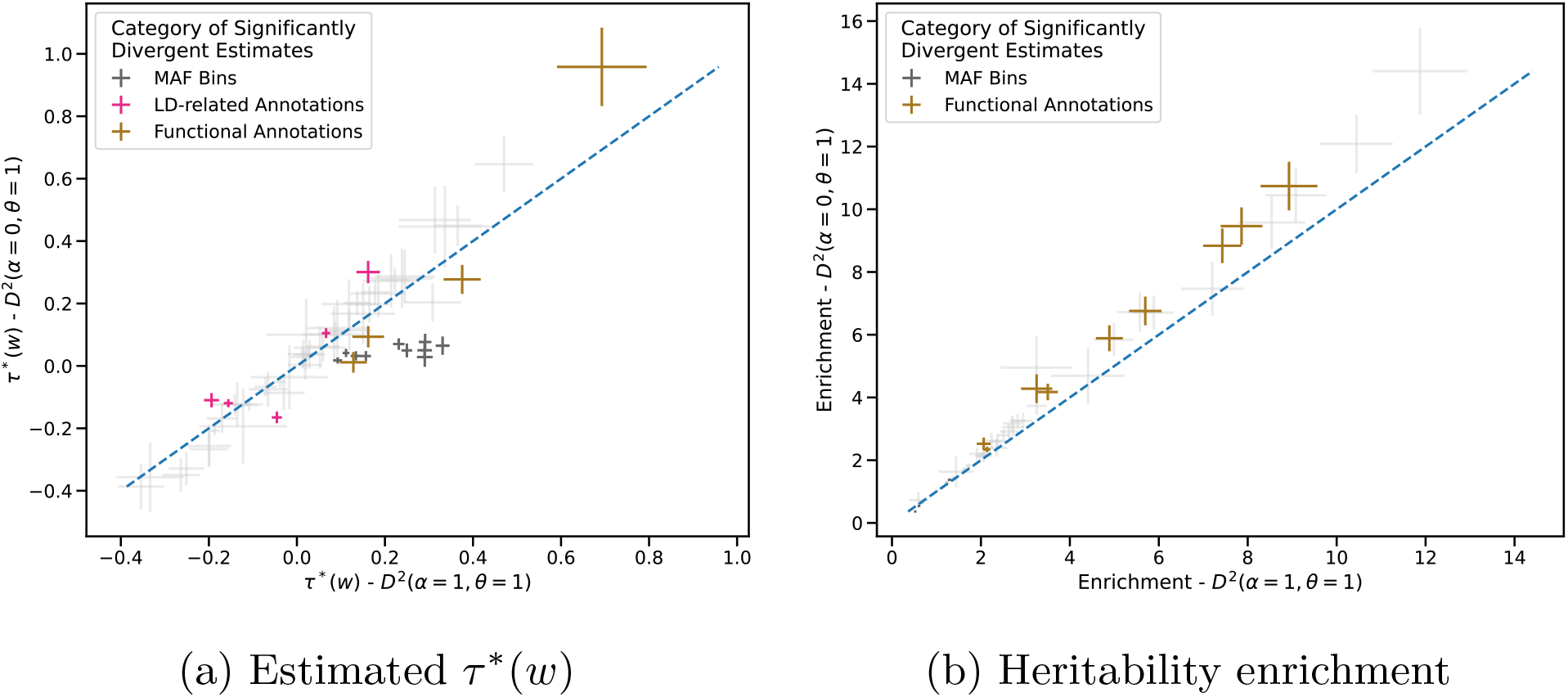
Comparison of meta-analyzed standardized coefficients and functional enrichment across different models of SNP heritability. (*α* = 1 on the x-axis and *α* = 0 on the y-axis). The estimates are meta-analyzed over 47 GWAS traits using a random-effects model. Estimates are shown with standard errors. **(a)** shows standardized coefficients *τ*^∗^(*w*) and **(b)** shows estimated heritability enrichment.

Figure 3(a) also has a slope higher than one, indicating that the choice *α* = 0 leads to stronger estimates of enrichment for highly enriched categories. We see a similar effect when examining heritability enrichment, a measure of enrichment commonly-used for binary annotations. Finucane et al., 2015 (Figure 3(b), Supplementary Table S9). Heritability enrichment is the ratio of the proportion of heritability explained by SNPs in a given functional category to the proportion of SNPs in that category. As expected, intermediate values of *α* produce less systematic shift in the estimates of enrichment for both statistics (Supplementary Figure 10).

To test the significance of the enrichment estimates, we compute *p*-values using the differential enrichment metric as defined by Hujoel et al., 2019 (see Appendix B). Even though *α* = 0 produce higher enrichment results for strongly enriched categories, two functional categories that are deemed highly significant under the *α* = 1 model fail to reach the significance threshold under the *α* = 0 model (Enhancer (Hoffman) and TSS (Hoffman), Supplementary Table S9). For example, Enhancer (Hoffman) has unadjusted p-values of 0.229 under *α* = 0 and 0.0005 under *α* = 1. The opposite effect is observed for the Weak Enhancer (Hoffman) functional category, where the *α* = 0 model reports it to be highly significant (Supplementary Table S9). These results again highlight the important role that the *α* parameter plays in the stratified LDSC framework, with different values of *α* potentially leading to different interpretations about the genetic architecture of complex traits.

### 3.4 The influence of *α* on models of SNP heritability

To explain the systematic shifts in enrichment as a function of *α* in the stratified models, here we empirically explore predicted mean squared effect size Var(*β*_*j*_) and corresponding association statistic 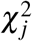 as a function of allele frequency under the LDSC models.

In the standard univariate model (*θ* = 0, *α* = 1), according to Equation 2, rare variants are predicted to have very large effect relative to common variants (Figure 4(a)). As a result, the model tends to strongly overestimate the chi-square statistics for rare variants, and underestimate them for common variants (Figure 4(b)). This effect is more pronounced for *α* = 1 than for any other choice of *α* (Figure 4(b)) and it helps explain the general trend of the standard univariate model producing downwardly biased estimates of SNP heritability (Speed et al., 2017): to avoid having large squared errors for rare variants in the regression, the model must underestimate the effect sizes of common variants, thereby underestimating the total heritability. This bias can be reduced by choosing a more biologically plausible value of *α*, or by using stratified regression.

**Figure 4.**
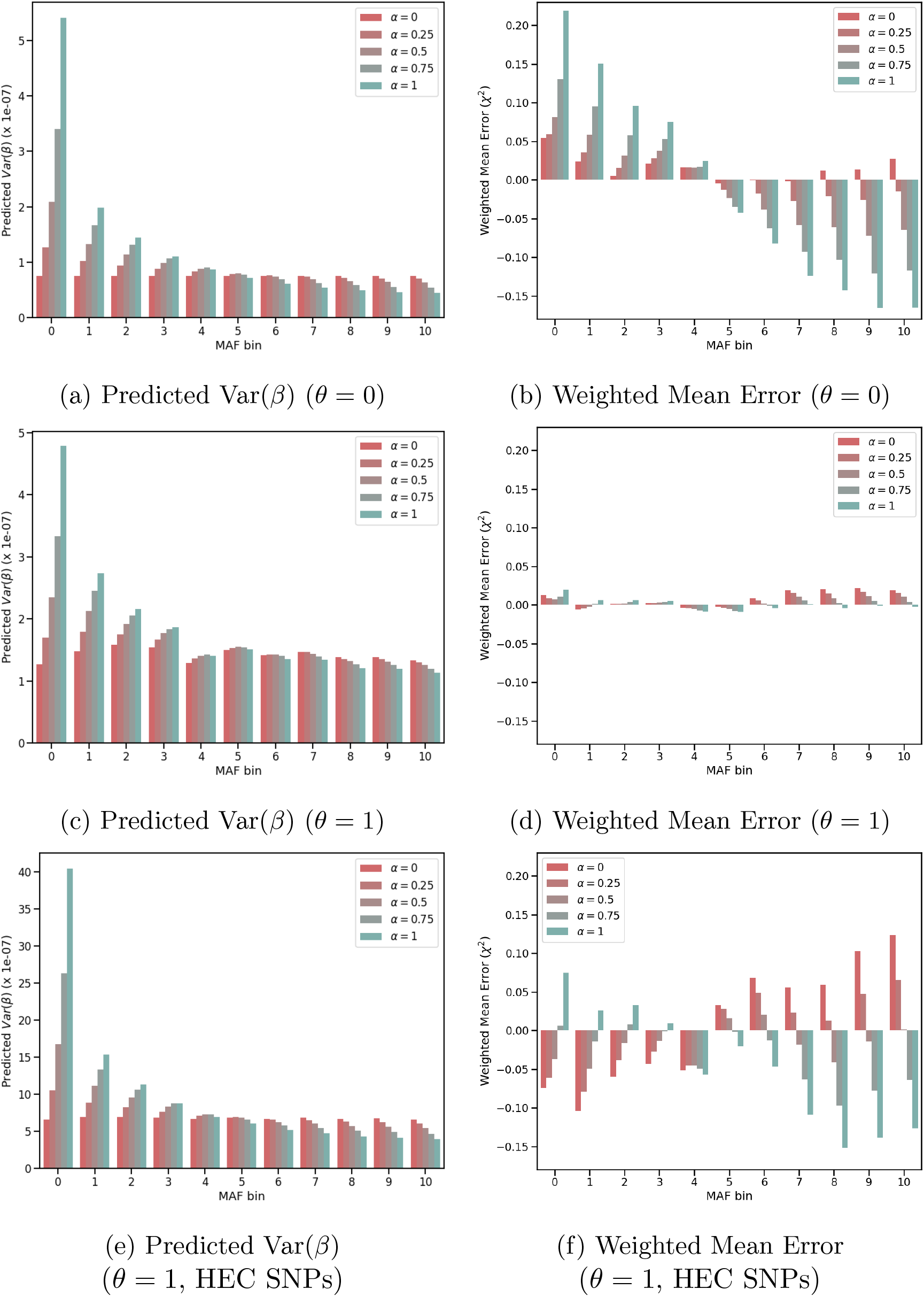
The predicted mean squared effect size and the mean error between predicted and observed *χ*^2^ for SNPs in different MAF bins with different values of *α*. Estimates are averaged across 47 GWAS traits. **(a, c)** show predicted mean squared effect size for SNPs under **(a)** univariate models and **(c)** stratified models. **(b, d)** show mean error between predicted and observed *χ*^2^ for SNPs under **(b)** univariate and stratified models. Panels **(e, f)** show predicted mean squared effect size and mean error for SNPs in the top 10 Highly Enriched Categories (HECs) under the stratified models.

We expect the stratified models to better describe the relationship between allele frequency and effect size, since these models have the freedom to assign arbitrary mean predicted effect sizes for each frequency bin. Indeed, all stratified models correctly overcome the bias in predicting the *χ*^2^ statistic (Figure 4(d)). However, they do so while predicting different effect sizes for rare and common variants (Figure 4(c)). Consequently, the weight of rare variants in the heritability of a trait can vary even in stratified models that agree on estimates of overall heritability. This, in turn, implies that the choice of *α* would lead to different proportions of heritability explained by each frequency bin and thus impact the predicted functional enrichment of variants in those bins (Figure 5(a)).

**Figure 5.**
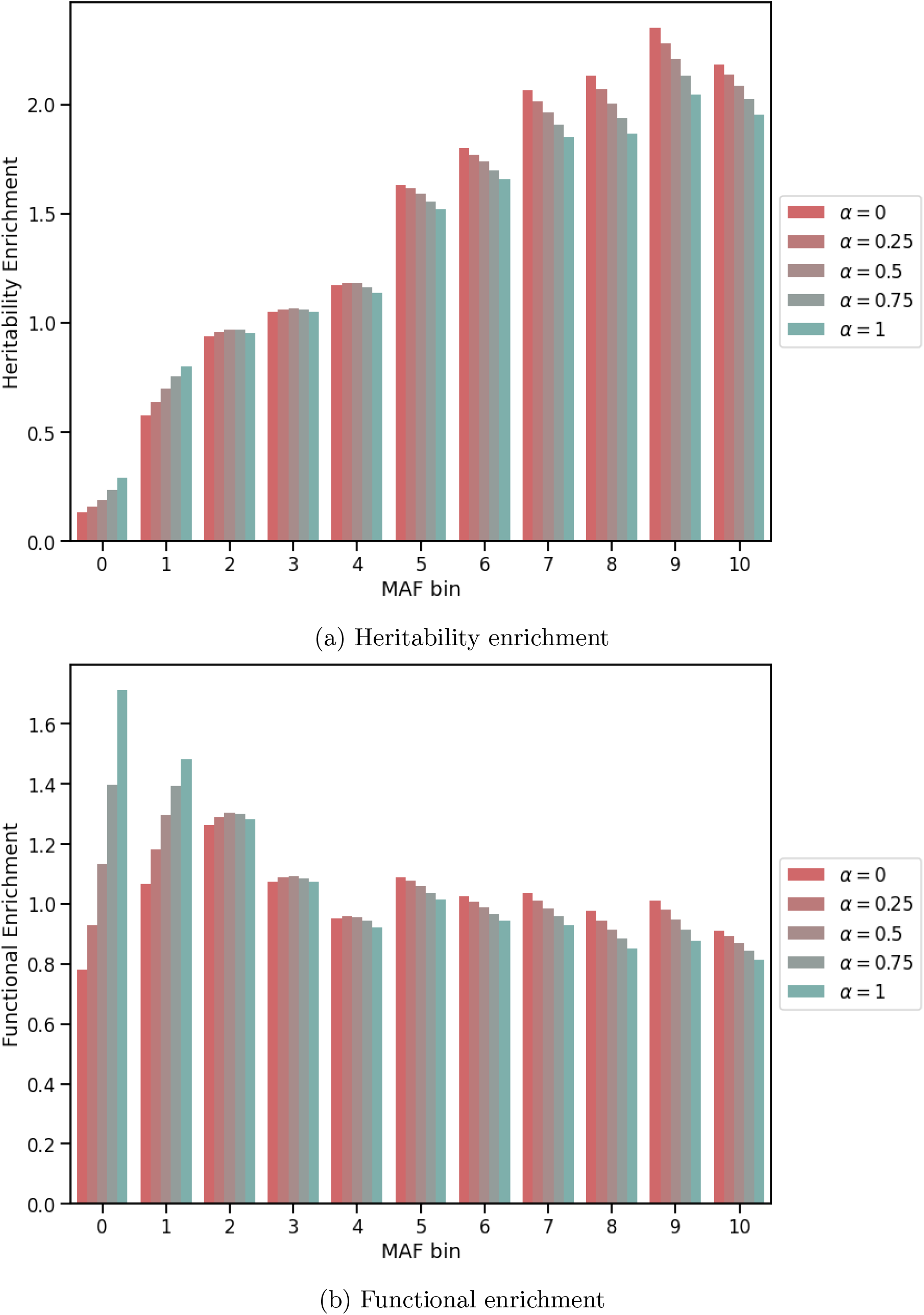
Estimates of enrichment for the MAF bins for stratified LDSC model with five different values of *α*. Estimates are averaged across 47 GWAS traits. **(a)** shows the predicted heritability enrichment metric for 11 MAF bins and **(b)** shows the functional enrichment metric for the same categories.

To explain the systematic differences in enrichment affecting highly enriched annotations, we now focus on the SNPs in the top 10 highly enriched functional categories (HECs, Supplementary Table S11), which comprise roughly 7% of all SNPs. The predicted mean effect size for rare functional variants is again very high for rare variants when *α* = 1 (Figure 4(e)). And, as in the univariate models, this results in correspondingly high weighted mean errors in estimating the *χ*^2^s for those SNPs (Figure 4(f)). This makes sense given the linear model for Var(*β*): for variants in strongly enriched categories, the annotation term overwhelms the frequency corrections and the distribution of effect sizes again follows the implicit assumptions induced by our choice of *α* (Figure 4(e)). Because the frequency bins fail to capture the frequency dependence for variants in highly enriched categories, the LDSC model for these variants is similar to a univariate model in that frequency effects are dictated by the choice of *α*. As in the univariate case, effects of rare variants are overestimated and, as argued above, the heritability contributed by these SNPs is underestimated. Figure 4 (b), (d), and (f) also show that in all cases, intermediate *α* produce less biased predicted chi-squared, as measures by the weighted mean error, than either *α* = 0 and *α* = 1.

Finally, the choice of *α* also has a large impact on the inferred architecture of complex traits as measured by the proportion of heritability attributed to common and rare variants. Figure 5(a) shows markedly different estimates of the heritability enrichment metric among rare variants. Figure 5(a) also shows that heritability enrichment should not be used as a proxy for functional enrichment: Even though common variants have lower per-allele effect size (e.g., Figure 4c), they are highly enriched for heritability relative to expectations under the (unrealistic) *α* = 1 model (Figure 5(a)). If the goal is to identify classes of variants with high per-allele effect size, we advocate for a different measure of functional enrichment, where contribution to heritability is measured relative to the expectation under a constant per-allele effect size (*α* = 0) (see Appendix B). Figure 5(b) shows the estimated functional enrichment across frequency bins, revealing the expected behaviour of slight depletion for common variants and slight enrichment for rare variants, with substantial differences depending on *α*. The 0th bin shows the largest difference across choices of *α*, and also an unexpected functional depletion for the rarest variants and *α* = 0. This 0th bin, which includes variants with allele frequency less than 5%, is often excluded from heritability estimates because of sensitivity to model assumptions and of other technical reasons (Bulik-Sullivan, Loh, et al., 2015).

## 4 Discussion

In this work, we proposed a reformulation of the LD Score Regression framework that replaces the *r*^2^ measure of LD with the *D*^2^ statistic, leveraging recently-proposed unbiased estimators (Ragsdale and Gravel, 2020). The reformulation highlighted the implicit assumptions about the relationship between a variant’s minor allele frequency and its effect size in commonly-used stratified LDSC models. This relationship, characterized by the parameter *α*, has been the subject of recent work on the frequency-dependent architecture of complex traits (Schoech et al., 2019; Speed et al., 2017; Speed et al., 2020; Zeng et al., 2018).

In the initial LDSC model, this parameter was set to *α* = 1 for mathematical convenience. Over the years, this assumption has been shown to be biologically implausible and to introduce biases in heritability estimates (Speed et al., 2017). These biases have been primarily addressed in the literature through stratified LDSC, giving the model more parameters and thus more flexibility to overcome the unrealistic assumption. Despite the additional parameters, we have shown that the choice of *α* still has a substantial effect on partitioned heritability metrics, such as standardized heritability coefficients and functional enrichment.

More recently, Speed et al., 2020 introduced a new heritability model, dubbed BLD-LDAK-alpha, that combines features of the Baseline LD and the LDAK models of SNP heritability and treats alpha as a free parameter that can be estimated from data. This is a promising approach that may alleviate the issues we report here, and the BLD-LDAK-alpha model results are generally consistent with our findings (Supplementary Figure 11). A notable difference between the two approaches is that the models presented here allow for a more flexible dependence of effect sizes on allele frequencies, both by conditioning on MAF bins and fitting the model along a coarse grid of alpha values, while the BLD-LDAK-alpha model allows for more systematic inference of the parametric dependence. The approach taken here allows us to investigate the effect of implicit assumptions on the commonly-used stratified LDSC model and to propose a minimal modification that largely resolves the issue, and to gain further insight into the interpretation of enrichment results.

These biases in partitioned heritability metrics exist because the choice of *α* = 1 implies rigid assumptions about the relationship between allele frequencies and effect sizes for variants in highly enriched categories. Under genome-wide empirical estimates of *α* = 0.38 (Schoech et al., 2019) and no annotation (*theta* = 0), Equation 2 predicts that the expected squared effect size of an uncommon variant at frequency *p* = 0.05 is 88% higher than a common variant at *p* = 0.5. Under the *α* = 1 model, that ratio is expected to be at 526%. Because the contributions of highly enriched functional annotations dominate the frequency corrections in Equation 2, this effect is still present for highly enriched variants in the model with annotations (*θ* = 1). These predictions are indeed borne out in the empirical results in Panels (a), (c), and (e) of Figure 4. Importantly, these panels show that conditioning on allele frequency bins is not sufficient to counteract the effect of setting *α* to an unrealistic value.

Even though such a synergistic effect is possible in theory, the analyses presented above and recent empirical estimates (Speed et al., 2020) suggest that values in the range *α* = 0.2 − 0.5 are much more plausible across all functional annotations. Even though uncertainty remains about the ‘best’ choice for *α*, we find much more modest differences in inference results if *α* is chosen within this range. Therefore, to avoid letting the best be the enemy of the good, we recommend using the genome-wide average *α* = 0.38 as a starting point for most stratified analyses.

## Appendix A Derivation of LD Score Regression with the *D*^2^ Statistic

Our re-formulation of the LD Score Regression model starts with a standard polygenic model where the phenotype of *N* individuals is assumed to linearly depend on *M* SNPs:

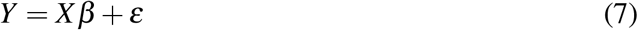

In this context, we assume that *Y* is a standardized vector of measured phenotypes and *X* is a *N* × *M* mean-centered (but not variance-normalized) genotype matrix. Following the derivation in Finucane et al., 2015, we express the Ordinary Least Squares (OLS) solution for the marginal statistic of SNP *j* as a function of *Y* :

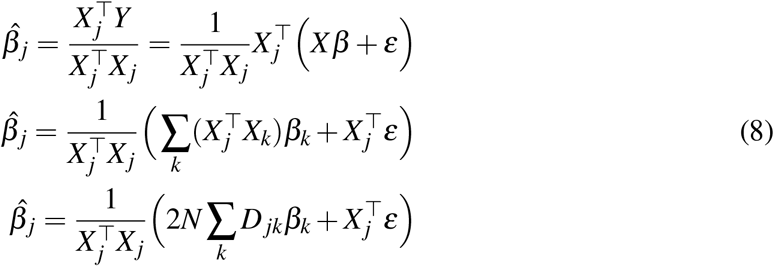

Where the third line follows from the definition of the *D* statistic 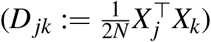. If we define the *χ*^2^ association statistic for SNP *j* as 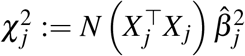 and take the expectation over all the random components, we obtain:

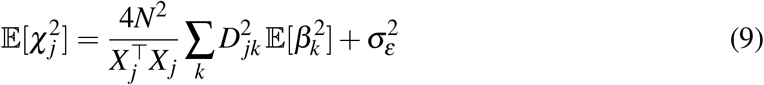

Under this model, the total narrow-sense SNP heritability is defined as 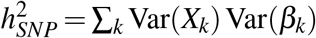. Given this, coupled with the assumption that 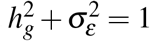, we obtain our general model for LD Score Regression with the *D*^2^ statistic:

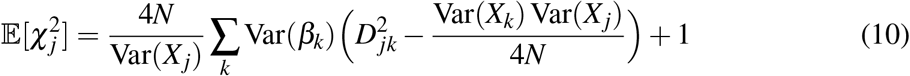

Assuming a large sample size for GWAS, this can be approximated by:

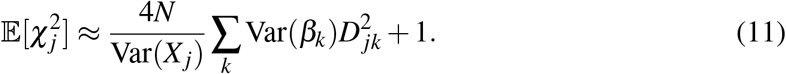

Finally, using the SNP heritability models outlined in Equation (2), the expression above can be equivalently formulated in terms of LD scores:

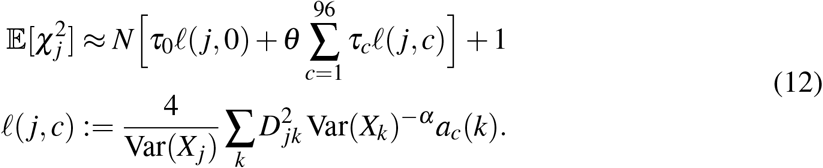

## Appendix B Definitions of SNP Heritability and Functional Enrichment

### B.1 SNP Heritability

SNP heritability quantifies the amount of phenotypic variance explained by a set of SNPs, genotyped or imputed (Yang, Zeng, Goddard, Wray, and Visscher, 2017). If we assume that the phenotype is standardized and the genotype matrix is mean-centered (but not variance-normalized), we can write our model of SNP heritability as:

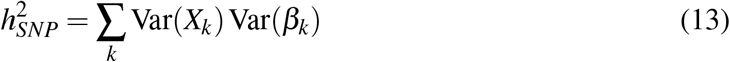

Using the above formulation, the models of SNP heritability outlined in Equation (2) can be expressed as:

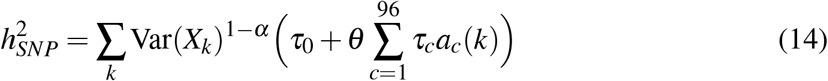

From the above expression, we can see that the relationship between partitioned heritability coefficients *τ*_*c*_ and the total SNP heritability 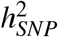 depends on *α*. For instance, in the univariate case (*θ* = 0), the coefficient *τ*_0_ is proportional to per-SNP heritability and is related to the total SNP heritability as

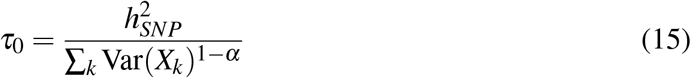

When *α* = 1, we recover the original LDSC formulation where the per-SNP heritability was assumed to be proportional to 1/*M* (Bulik-Sullivan, Loh, et al., 2015). On the other hand, when *α* = 0 the per-SNP heritability is inversely proportional to the average allele frequency variance across all SNPs.

### B.2 Heritability and Functional Enrichment

Heritability enrichment is defined as the ratio of the proportion of heritability explained by SNPs in a given category *c* relative to the proportion of SNPs in that category (Finucane et al., 2015):

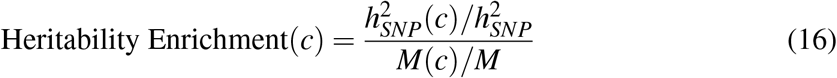

Where 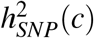 is the heritability explained by SNPs in category *c* and *M*(*c*) is the number of SNPs in that category. To test for significance of the functional enrichment metric, we use the differential enrichment metric as defined in Hujoel et al., 2019:

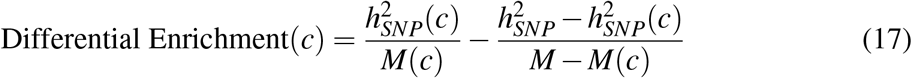

As outlined in the main text (Section 2.5), a better measure of functional enrichment would be measuring contributions to heritability relative to expectation under a constant per-allele effect size (*α* = 0) model. This implies that instead of dividing by the proportion of SNPs, we divide by the proportion of genotypic variances:

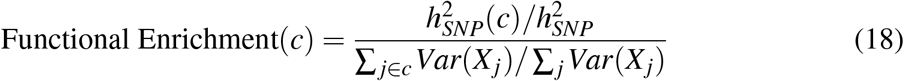

## Appendix C Analysis Procedures and Evaluation Metrics

### C.1 Computing and Comparing LD Scores from 1000 Genomes Data

To conduct the analyses discussed in this paper, we computed LD scores using three different estimators of LD (naive *r*^2^, corrected *r*^2^ (Bulik-Sullivan, Loh, et al., 2015; Yin and Fan, 2001) and *D*^2^ (Ragsdale and Gravel, 2020)) for *α* ∈ {0.0, 0.25, 0.5, 0.75, 1.0} and for 3 of the super populations in Phase III of the 1000 Genomes Project (Africans, Asians, and Europeans) (Altshuler et al., 2010; The 1000 Genomes Project Consortium, 2012), excluding the sets of closely related individuals identified in Gazal, Sahbatou, Babron, Genin, and Leutenegger, 2015. We followed the same quality control procedures as in the original LDSC software documentation (Bulik-Sullivan, Loh, et al., 2015) (e.g. discarding singleton SNPs). All of these quality control steps were run with `PLINK v1.9` (Purcell et al., 2007).

To qualitatively compare LD Scores from the univariate model with different values of *α*, we normalized them such that the slope of the univariate regression becomes 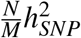 instead of *Nτ*_0_. Given our definition of *τ*_0_ in terms of 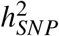 in Equation (15), we see that this can be achieved by dividing the LD score for each model by the following quantity:

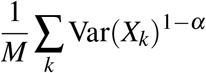

By construction, when *α* = 1, as in the standard LDSC models, the normalization factor is 1. When *α* = 0, the normalization factor becomes the average genome-wide variance in allele frequency.

### C.2 Parameter Estimation and Meta-analysis

To estimate the parameters of the various heritability models, we used the Iteratively re-weighted least squares (IRLS) jackknife estimator from Bulik-Sullivan, Loh, et al., 2015 with some modifications to account for differences in the heritability models. Primarily, since our general model starts with an unnormalized genotype matrix, the total SNP heritability is given by:

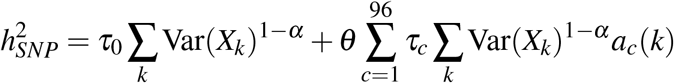

Once the coefficients 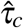 have been estimated by the `ldsc` software, to obtain the total heritability, we have to multiply them by different factors. For example, in the case of the *α* = 1 models, we multiply by the sum of the annotations ∑ _*j*_ *a*_*c*_(*k*), whereas for the *α* = 0 models, we multiply by the sum of the annotations weighted by MAF: ∑_*k*_ Var(*X*_*k*_)*a*_*c*_(*k*).

To meta-analyze the coefficients as well as measures of functional enrichment across the 47 GWAS traits, we used a random effects model as implemented in the `R` package `meta` (Balduzzi, Rücker, and Schwarzer, 2019).

### C.3 Empirical Evaluation of SNP heritability models

To empirically evaluate the stratified SNP heritability models as in Section 2.5, we fit each model to the summary statistics from 47 GWAS traits (Supplementary Table S1) using the `ldsc` software (Bulik-Sullivan, Loh, et al., 2015) with standard configurations. For each trait and model, we obtained the mean parameter estimates 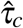 and used them to compute the predicted 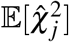 for all SNPs that were used in the regression. Then, for each MAF bin, we computed the weighted mean error using the following equation:

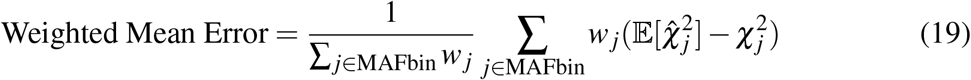

Where the 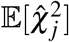 is the predicted association statistic under the model and 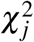 is the observed statistic. The weight *w*_*j*_ is simply the LD score weight that we employed in the regression Bulik-Sullivan, Loh, et al., 2015. This strategy of weighing the performance statistics by the LD score weights has been explored in previous work Speed et al., 2020. Since, in our case, each model has its own set of LD score weights that match its *α* value, here we used the LD score weights for the *α* = 1 model throughout. The weighted mean errors per MAF bin that we report in the main text are averaged across the 47 GWAS traits.

## Supplemental Data

Supplemental Data include ten figures (Supplementary Figures S1-10) and eleven tables (Supplementary Tables S1-11).

## Acknowledgements

We thank Chris Gignoux and members of the Gravel Lab for useful discussions.

## Web Resources

Baseline-LD model version 2.2,

data.broadinstitute.org/alkesgroup/LDSCORE/

Summary statistics for 47 GWAS traits,

data.broadinstitute.org/alkesgroup/LDSCORE/independent_sumstats/

## Data and Code Availability

Code to compute the LD scores with the unbiased estimator of *D*^2^ and carry out the analyses discussed in this paper is available on github: https://github.com/shz9/unbiased-ldsc.

